# Carbon starvation of *Mycobacterium abscessus* induces a non-replicating state with extensive proteomic remodeling

**DOI:** 10.64898/2026.05.05.723019

**Authors:** Kaylyn L. Devlin, Gyanu Lamichhane, William C. Nelson, Vivian S. Lin, Kimberly E. Beatty

## Abstract

*Mycobacterium abscessus* (*Mab*) is an opportunistic pathogen that can cause chronic, debilitating lung disease. *Mab* is intrinsically resistant to most antibiotics, making *Mab* infections challenging to manage and frequently incurable. During infection, *Mab* adapts to survive various stresses, including hypoxia and nutrient starvation. *In vitro*, these conditions drive *Mab* into a drug-tolerant, non-replicating state. Changes in the *Mab* proteome that result from entering a non-replicating state have been minimally described despite the clinical importance of this physiological state. Using *Mab* reference strain ATCC 19977, we collected proteomic data comparing replicating to non-replicating states using a carbon starvation (CS) model of persistence. We identified 2251 proteins overall (46% proteome coverage), and 17% of these proteins were found in only one of the two conditions. A third of identified proteins were significantly changed in abundance, indicating an extensive proteomic response to CS. The response regulator DosR and many DosRS responsive proteins were significantly more abundant under CS, suggesting that the DosRS stress response regulator plays a key role in CS-induced *Mab* persistence. Many aspects of cell wall biosynthesis were changed, including changes in glycolipid abundance under CS. Proteins involved in other key cellular processes such as secretion, oxidative phosphorylation, and nutrient metabolism were altered under CS. The proteomic analysis presented provides new insights and clarity into how the *Mab* proteome is regulated during non-replicating persistence, a key consideration for understanding *Mab* pathophysiology.

## Introduction

*Mycobacterium abscessus* (*Mab*) is a non-tuberculosis mycobacteria (NTM) that is ubiquitous within water- and soil-based environmental niches. *Mab* infections are most commonly acquired by inhalation and therefore establish within the lung, although *Mab* can also infect the skin and soft tissues following trauma^1^. People with chronic lung conditions, such as bronchiectasis, chronic obstructive pulmonary disease, or cystic fibrosis, are most vulnerable to *Mab*-related disease. Infections are incredibly hard to cure due to intrinsic drug resistence, a fact that gives *Mab* the infamous designation as a “clinical nightmare”^2^.

A hallmark of mycobacterial drug resistance is the ability of the pathogen to enter a non-replicating persistent state in response to host conditions, such as low pH, hypoxia, nutrient deprivation, and other factors^3^. *Mab* is exposed to these stresses in the host within the granuloma^4,5^ and through biofilm formation^6,7^. Induction of a non-replicating state *in vitro* by modeling host conditions has demonstrated that persistent *Mab* are phenotypically drug tolerant^8–10^, a factor that contributes to treatment failure. Efficacy testing of novel drugs against *Mab* is now encouraged to include persister assays to better predict clinical outcomes^3^.

Despite the acknowledged clinical importance of persistent *Mab*, little is known about the proteomic characteristics of non-replicating state. A thorough understanding of the pathophysiological adaptations of *Mab* to changing environments could improve treatment strategies or identify novel therapeutic targets. Several studies have investigated transcriptional changes in *Mab* persisters induced through exposure to hypoxia^11,12^, nutrient starvation^13^, and growth in artificial sputum^12^. However, transcription and translation are often decoupled^13^, making the proteome more reflective of physiology than the transcriptome. To date, proteomic regulation by *Mab* persisters has been minimally described using models based on hypoxia^14^, complete nutrient starvation, or potassium depletion^13^.

In the current work, we sought to expand the understanding of *Mab* persistence and related proteomic adaptations. We used a new model of non-replicating persistence based on previously described nutrient starvation models^8–10,15,16^, which restricts carbon while maintaining inorganic nutrients. Our studies were completed in the reference strain, *M. abscessus* subspecies *abscessus* (ATCC 19977). We compared the proteome of carbon starved, non-replicating *Mab* to actively replicating *Mab*. As described herein, we identified profound proteomic changes that occur in response to carbon starvation, spanning key mycobacterial functions such as secretion, energy metabolism, virulence, and cell envelope biosynthesis and composition.

## Results and Discussion

### A model for non-replicating persistence

Most *in vitro* models for *Mab* persistence have been adapted from those optimized for *Mycobacterium tuberculosis* (*Mtb*), such as the Wayne model^17^ of hypoxia or the Betts model^18^ of nutrient starvation. For example, Berube *et al*.^8^ first adapted the Betts model for *Mab*, where bacteria were placed in phosphate-buffered saline (PBS) supplemented with tyloxapol for four days. They found that nutrient-starved *Mab* retained susceptibility to amikacin but was resistant to most other drugs. Yam and coworkers^9^ also used a PBS-based nutrient starvation model, reporting suspension of growth and maintenance of viability over ten days, along with high tolerance to a range of antibiotics in these non-replicating persisters.

However, we observed an initial decrease in *Mab* colony-forming units (CFUs) when cultured in PBS^19^. As a result, we based our non-replicating *Mab* model on the Hung model^15^ of carbon starvation, which has been used in numerous studies of *Mtb*^16,20–22^. *Mab* was cultured for three days in Middlebrook 7H9 broth supplemented with tyloxapol only (carbon starvation; CS), constituting starvation of carbon with the maintenance of inorganic compounds. CS halted growth and maintained viability, without the initial apparent cell death observed in PBS-only culture. We therefore used the CS induced non-replicating state as a model of *Mab* persistence for our proteomic analyses.

### Global proteomics analysis

Global proteomic profiles were generated for *Mab* grown under CS for 72 hrs and for actively growing *Mab* from nutrient-complete cultures (REP) using mass spectrometry-based proteomics. To ensure rigor, we only considered proteins that were identified in three or more of the six biological replicates of either group (REP or CS). Overall, we identified 2251 proteins out of the 4949 annotated in *Mab* (45% proteome coverage; **Table S1**). A similar number of proteins was identified in REP (2046) and CS (1956) conditions, and most proteins (1751) were identified in both (**Figure 1**).

**Figure 1.**
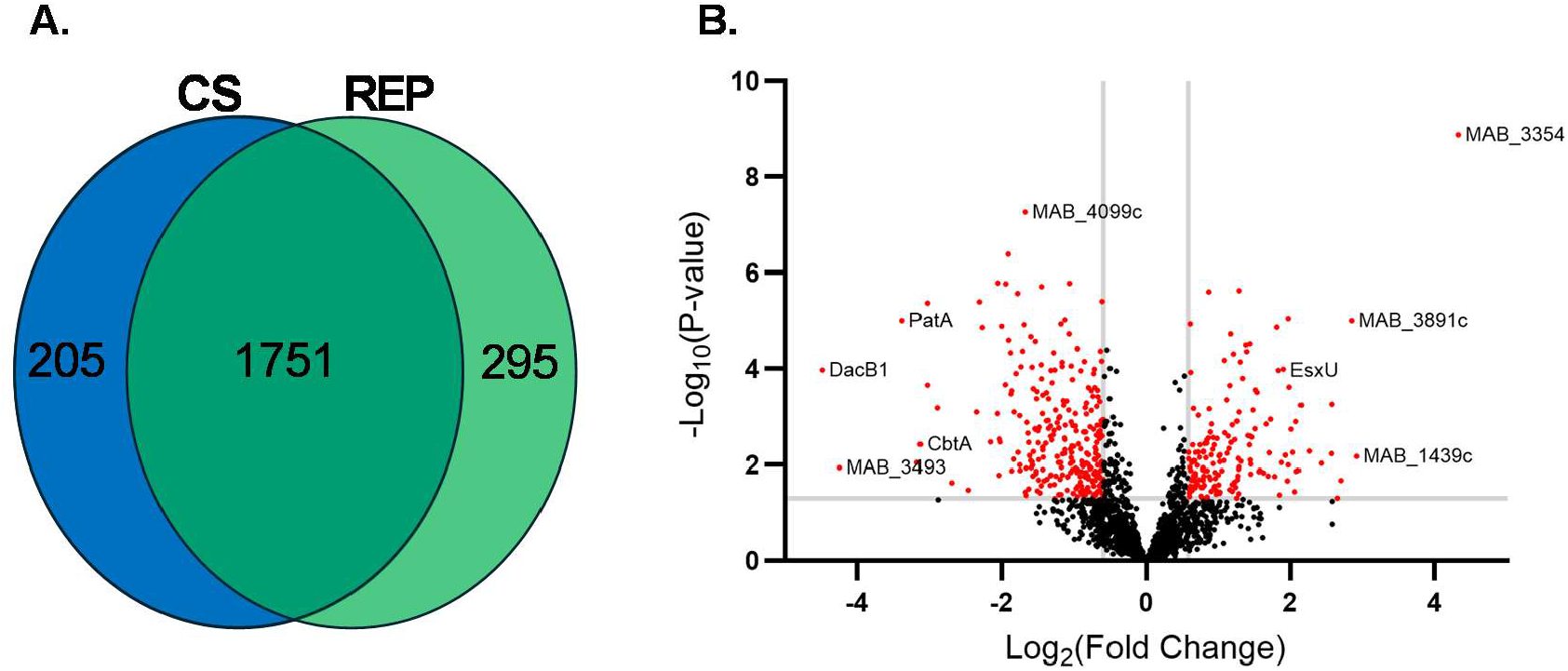
Comparison of proteins identified in CS versus REP conditions. **A**. Venn diagram of proteins identified in n ≥ 3 samples in CS or REP conditions. **B**. Volcano plot of proteins found in CS and REP conditions. Red dots highlight proteins significantly different in abundance between groups with p ≤ 0.05 (ANOVA) and a log_2_Fold Change (FC) in CS relative to REP ≥ 0.585 (more abundant) or ≤ −0.585 (less abundant). Proteins identified in only one group (n ≥ 3 in group A and n ≤ 1 in group B) were not included in the volcano plot due to the inability to calculate a p-value.

We used spectral intensities as an estimate of protein abundance to identify differentially-expressed proteins (DEPs) between growth conditions. We defined DEPs as proteins changed in either abundance (ANOVA, p ≤ 0.05, >1.5-fold, **Figure 1B**) or presence (G-test, p ≤ 0.05) (**Table S2**). There were 288 DEPs in CS, with 125 present only in CS (n ≤ 1 in REP). Conversely, there were 463 DEPs in REP, with 194 present only in REP (n ≤ 1 in CS). Overall, 33% of proteins in our dataset were significantly changed in response to CS, indicating substantial proteomic regulation by persistent *Mab*.

Functional classification of DEPs is challenging because the *Mab* proteome remains poorly annotated and 40% of proteins have no known function. We used *Mtb* orthology to enrich *Mab* protein classification. A multistep analysis was performed to identify orthologs between *Mab* 19977 and *Mtb* H37Rv (**Table S3**). Integration of reciprocal best hits, OrthoFinder^23^, Markov clustering, MMSEQS2 clustering^24^, and consideration of gene synteny were performed to, optimally, identify one-to-one gene mappings across organisms. In agreement with similar analyses^25–27^, less than 50% of *Mab* genes (2566) were identified to have a high-confidence ortholog in *Mtb*. However, 2334 genes were shared between the two species (**Figure S1**).

Using UniProt^28^ annotations and *Mtb* orthology, we assessed specific CS-related proteomic changes across key cellular and pathogenic processes including intra- and extracellular signaling, cell wall biosynthesis, and energy and nutrient metabolism (**Table S4**).

### Two-component signaling systems and DosRS

Mycobacterial two-component signal transduction systems (TCSS) regulate major cellular processes such as growth, persistence, and virulence in response to environmental signals^29^. Those signals can include low pH, oxidative stress, metals, nutrients, antibiotics, and other factors. TCSS are comprised of a pair of proteins including a periplasm-facing histidine sensor kinase that confers a signal to a cytoplasmic response regulator (**Figure 2**). Activated response regulators bind DNA to induce transcriptional changes.

**Figure 2.**
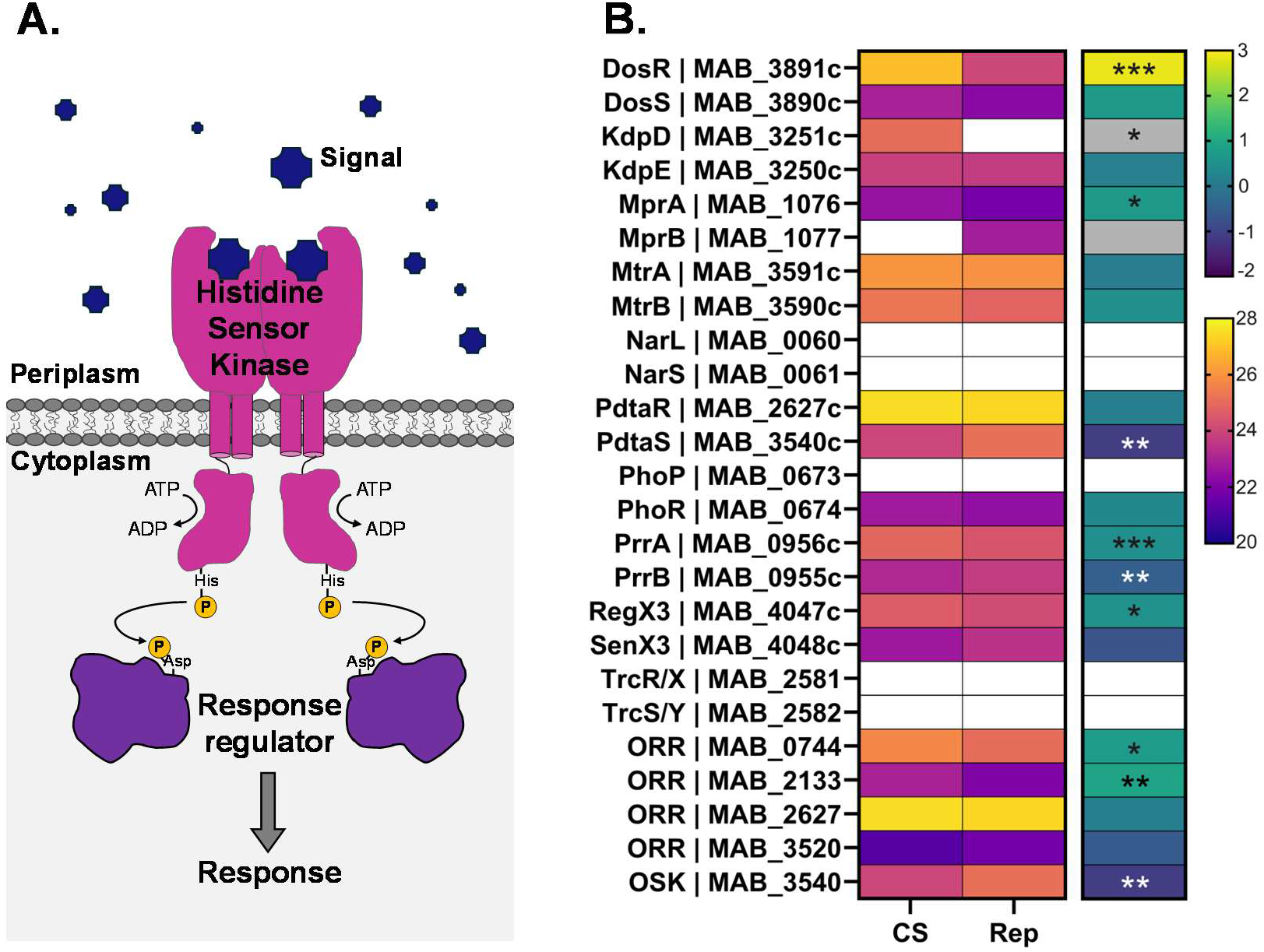
Two-component signaling systems are altered by carbon starvation. **A**. Schematic of a two-component signaling system, which uses a histidine sensor kinase and a response regulator to regulate gene expression in response to a signal from the environment. **B**. Heat map of mean log2 intensities (left map, warm scale) and corresponding log2 fold-change (right map, cool scale) of TCSS proteins in CS versus REP (n = 6) groups. A grey box indicates a protein was present in only one group (Group A: n ≥ 3, Group B: n ≤ 1). A white box indicates absence of value. Asterisks denote significance of the difference in presence (g-test) or mean intensities (ANOVA) between CS and REP (*: P ≤ 0.05, **: P ≤ 0.01, ***: P ≤ 0.001). ORR: orphan response regulator. OSK: orphan sensor kinase.

*Mab* encodes at least ten TCSS and five orphan components (regulators or kinases). We observed significant changes in abundance of nearly half of these proteins in response to CS (**Figure 2B**). The largest change was in DosRS. Prior studies showed that DosRS is required for growth, survival, and adaptation to hypoxia^11,30^. We found that the response regulator, DosR (MAB_3891c), was 7.2-fold more abundant in CS samples compared to REP (p = 1.0 × 10^−7^). The abundance of the sensor histidine kinase DosS (MAB_3890c) was comparable between CS and REP. Our findings are in accordance with other persistence and stress models, as transcripts of these regulators were found to be elevated in response to hypoxia (*dosR* and *dosS*)^11,12^ and intracellular survival within macrophages (*dosR* only)^31^. However, a recent study looking at *Mab* proteomic changes in response to total nutrient starvation did not find DosR or DosS altered at the protein level compared to active replication^13^. Therefore, our results provide the first evidence that DosRS is used by *Mab* to respond to carbon starvation.

We compared our dataset to DosRS-regulated genes identified as transcriptionally up-regulated in hypoxia. We limited our comparison to the ten most induced DosRS-regulated genes identified by either Simcox *et al*.^11^ or Belardinelli *et al*.^30^, for a total of 11 proteins (**Table S4**). Eight were identified in our dataset, seven of which were significantly more abundant in CS (> 4-fold; p < 0.0002). These proteins included a fatty acid desaturase DesA (MAB_3354), adenylate kinase (MAB_3902c), NADPH nitroreductase (MAB_3903), cytochrome c oxidase sub-unit (MAB_1042c), two universal stress protein A (UspA) domain-containing proteins (MAB_3904 and MAB_2489), and an unannotated protein (MAB_3937). The remaining identified protein within the comparison set, a nitric oxide dioxygenase (MAB_3133c), was only found in REP (p = 0.071). Overall, our proteomic data support that DosRS serves as a key regulatory mechanism for responding to carbon starvation, mirroring many of the responses observed under hypoxia.

There were changes in several uncharacterized *Mab* TCSS between CS and REP samples. We identified the sensor kinase KpdD (MAB_3251c) in CS alone (p = 0.023), while its paired response regulator KpdE (MAB_3250c) was unchanged between CS and REP. The orthologous *Mtb* TCSS is induced under nutrient stress^18^. PdtaR/S is uncharacterized in *Mab*, but we found that the sensor kinase PdtaS (MAB_3540c) was 2.2-fold less abundant in CS (p = 0.0058). There were two recent reports characterizing *Mtb*’s PdtaR/S. One found that it plays a role in virulence and persistence^32^. A second report found that PdtaR/S senses heme and regulates oxidative stress responses^33^. Next, SenX3/RegX3 regulates virulence in *Mtb*; the *Mab* RegX3 (MAB_4047c) response regulator was slightly, but significantly, more abundant in CS (1.4-fold; p = 0.015). The function of MprA/B in *Mab* is unknown, but these proteins are necessary for persistence in *Mtb*^34,35^. MprA was 1.6-fold more abundant in CS (p = 0.016), although its cognate kinase MprB was only identified in REP.

TCSS are under active investigation in *Mab* with several papers published since 2023. For example, MtrA/B was recently characterized to impact drug resistance, virulence, and cell division^36^. We identified the MtrA/B, but it was not significantly changed between REP and CS. Also recently characterized are the orphan response regulators GlnR (MAB_0744) and NnaR (MAB_3520c)^37^, which mediate nitrogen metabolism. GlnR, which also facilitates biofilm formation^38^, was 1.6-fold more abundant in CS compared to REP (p = 0.026), while NnaR was not significantly different. Lastly, PhoP/R mediates the response to low pH^39^; PhoR was identified but unchanged.

### Secretion systems

Type VII secretion systems (T7SS) are used by mycobacteria to export proteins and nutrients across the cell envelope (**Figure 3**). T7SS are linked to pathogenesis, immune system evasion, and survival in the host^40,41^. While *Mtb* encodes five T7SS, *Mab* encodes only two: Esx3 and Esx4. Esx3 (MAB_2224c-MAB_2234c) plays a role in growth, persistence, and pathological responses during infection^12,42^. We identified five components of the Esx3 core complex, and EccC3 (MAB_2232c) was 2.6-fold more abundant in REP than CS (p = 0.0012) (**Figure 3B**). We identified an Esx3 substrate, EsxG (MAB_2229c), only in REP samples. Both EccC3 and EsxG were recently found to be reduced at the protein level in response to total nutrient starvation, similar to our findings^13^.

**Figure 3.**
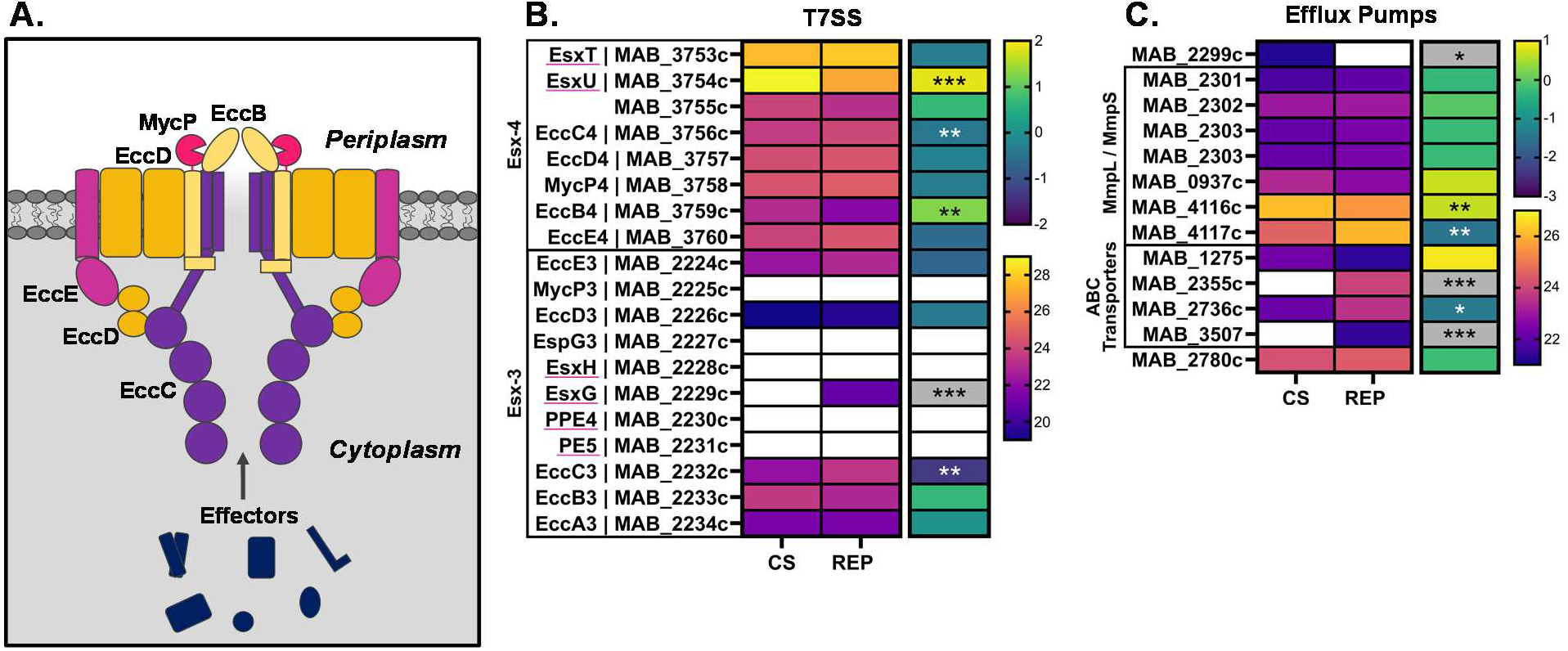
Organization and regulation of *Mab* secretion systems under CS. **A**. Functional organization of mycobacterial T7SS pore-forming components. **B**., **C**. Heat maps of mean log2 intensities (left map, warm scale) and corresponding log2 fold-change (right map, cool scale) of T7SS proteins (**B**.) and efflux pumps (**C**.) in CS versus REP (n = 6) groups. Scale bars are applicable across all corresponding maps. A grey box indicates a protein was present in only one group (Group A: n ≥ 3, Group B: n ≤ 1). A white box indicates absence of value. Asterisks denote significance of the difference in presence (g-test) or mean intensities (ANOVA) between CS and REP (*: P ≤ 0.05, **: P ≤ 0.01, ***: P ≤ 0.001). Underlined proteins in (**B**.) are substrates of Esx3 or Esx4.

Esx4 contributes to intracellular survival, blocking phagosomal acidification and inducing phagosomal rupture during infection^43^. All Esx4 components (MAB_3753c-MAB_3760) were identified in both conditions (**Figure 3B**). We found that a key structural component, EccB4 (MAB_3759c), was 2.4-fold more abundant in CS (p = 0.0013). Additionally, Esx4’s substrate EsxU (MAB_3754c) was 3.6-fold more abundant in CS (p = 0.00011). A recent study showed that heterodimeric EsxU/T is up-regulated during infection, where it is involved in phagosomal membrane rupture and reducing virulence^44^. Taken together, our data suggest that Esx4 may have an important role in *Mab* under conditions of CS, while Esx3 may be more relevant during active growth.

The induction of efflux pumps is hypothesized to contribute to the emergence of drug resistance in mycobacterial pathogens^45^. Several recent studies demonstrated that *Mab* upregulates efflux pumps in response to antibiotic treatment^46–50^. We examined our data and found some interesting changes in both efflux pumps and their transcriptional regulators (**Figure 3C**). For example, MAB_2299c, a transcriptional regulator associated with clofazimine and bedaquiline resistance^49^, was identified only in CS. The corresponding membrane proteins, MAB_2301 and MAB_2302, were unchanged. An efflux pump linked to macrolide resistance^48^, MAB_2355c, was found only in REP. Lastly, the ABC transporter MAB_3507 was found only in REP.

### The mycobacterial cell envelope

#### PG biosynthesis and degradation

Peptidoglycan (PG) is a massive polymer that encapsulates bacterial cells and serves as their exoskeleton. In mycobacteria, it is comprised of a glycan backbone of N-acetylglucosamine and N-acetylmuramic acid attached to peptide stems, which are highly cross-linked. PG biosynthesis commences in the cytoplasm with synthesis of UDP-N-acetylglucosamine attached to the peptide stem. These steps are catalyzed by GlmS (MAB_3743), GlmM (MAB_3750c), GlmU (MAB_1148c), and MurA/B/C/D/E/F/G/I. NamH (MAB_0156c), alanine racemase (Alr; MAB_3739c) and D-Ala-D-Ala ligase (DdlA; MAB_3286c) also play a role^51^. All of these proteins were identified in both CS and REP (**Figure 4A**). Two were more abundant in REP than CS (GlmU and MurA) and MurI was only identified in REP conditions. Rojony *et al*.^14^ reported that MurC/D/E/F/G/X were upregulated in hypoxia and biofilm conditions, but we found no significant differences.

**Figure 4.**
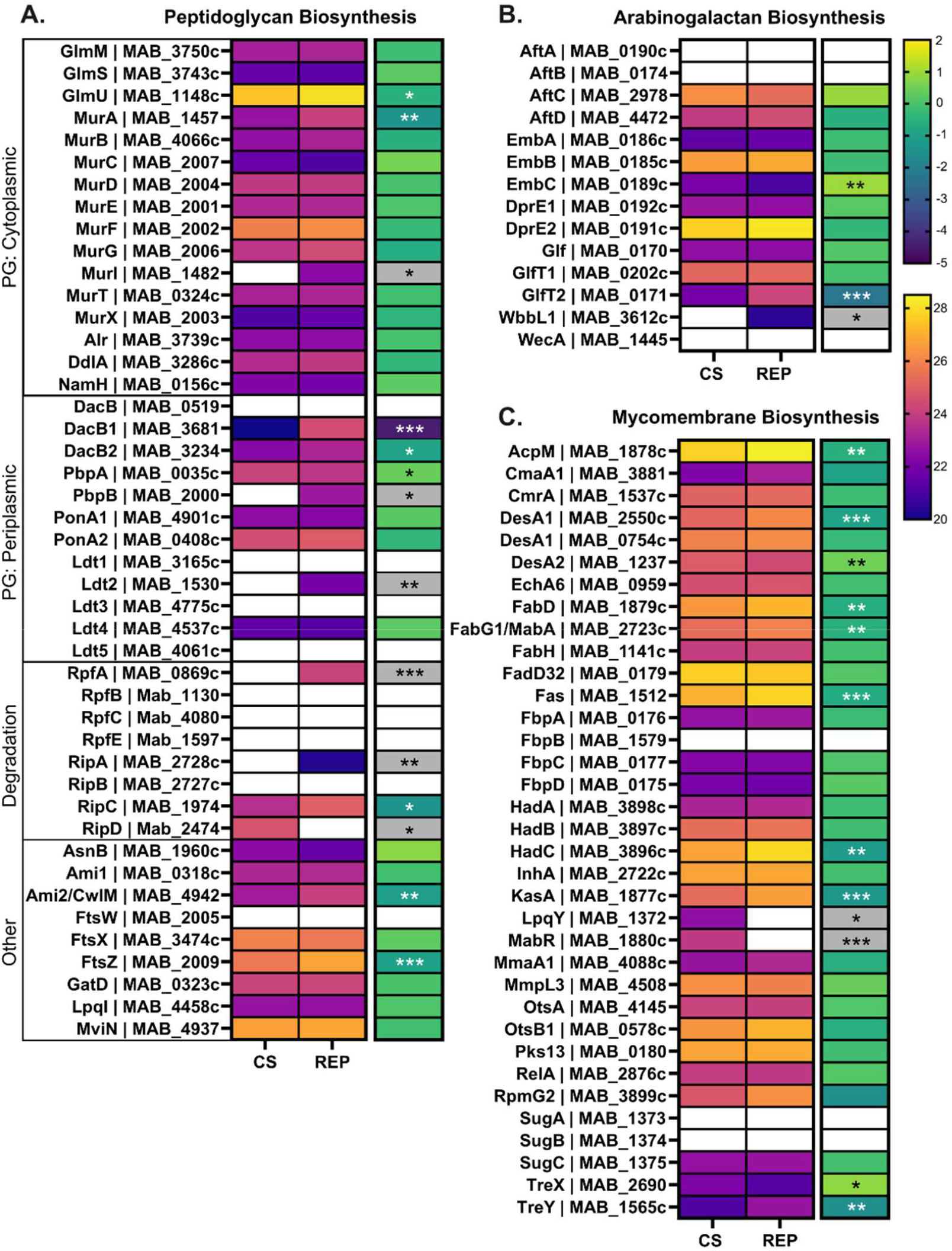
Regulation of cell envelope biosynthesis under CS. Heat maps of mean log2 intensities (left map, warm scale) and corresponding log2 fold-change (right map, cool scale) in CS versus REP (n = 6) groups of proteins involved in peptidoglycan (**A**), arabinogalactan (**B**), and mycomembrane (**C**) biosynthesis. Scale bars are applicable across all corresponding maps. A grey box indicates a protein was present in only one group (Group A: n ≥ 3, Group B: n ≤ 1). A white box indicates absence of value. Asterisks denote significance of the difference in presence (g-test) or mean intensities (ANOVA) between CS and REP (*: P ≤ 0.05, **: P ≤ 0.01, ***: P ≤ 0.001). In panel **C**, DesA1 (MAB_3354) was excluded from the heat map due to the large fold change skewing color mapping.

The remaining biosynthetic steps take place in the periplasm, where carboxypeptidases trim the peptide stems and transpeptidases cross-link them. *Mab*’s PG contains two distinct types of cross-links. The first are classically described D-Ala-D-Ala (4⟶3) cross-links, which are formed by D,D-transpeptidases (DDTs). These enzymes are also called penicillin-binding proteins. We identified four DDTs in our dataset: PbpA (MAB_0035c), PbpB (MAB_2000), PonA1 (MAB_4901c), and PonA2 (MAB_0408c). PbpA was slightly more abundant in CS (1.3-fold, p = 0.042) and PbpB, an essential enzyme^26^, was only identified in REP samples. Mycobacterial PG also has mDap-D-Ala (3⟶3) “non-classical” cross-links. The latter predominate in *Mab* and are formed by L,D-transpeptidases (LDTs) using a tetrapeptide substrate made by carboxypeptidases^52^. The carboxypeptidases DacB1 (MAB_3681) and DacB2 (MAB_3234) were identified in our dataset. Both were substantially more abundant in REP samples, with DacB1 being among the most CS-responsive proteins at 22.3-fold less abundant in CS than REP (p = 0.00011). We found Ldt4 (MAB_4537c) in both CS and REP, while Ldt2 (MAB_1530) was only found in REP samples. Overall, our data demonstrate that many enzymes associated with PG biosynthesis are less abundant in CS and distinct PG-modifying enzymes are used in replicating versus non-replicating states. We recently described results from activity-based protein profiling in REP and CS *Mab*, where β-lactam antibiotic targets were identified using a biotinylated meropenem^19^. We determined that carboxypeptidases, DDTs, and LDTs were active in both growth conditions, suggesting that these proteins remain druggable in CS despite reduced abundance.

There are various PG degrading enzymes encoded by *Mab*, including the resuscitation promotion factors (Rpfs A-E) and endopeptidases (RipA-D). In *Mtb*, RpfA is required for recovery of growth after dormancy^53^ and RipA is localized to the septa of dividing cells^54^. Our data indicate that these degraders are regulated in response to CS. We found RipD in CS alone, while RpfA, RipA, and RipC were more abundant or found only in REP. Grigorov *et al*.^13^ reported similar results, with RpfA and RipA proteins reduced in total nutrient starved *Mab*. The absences of RpfA and RipA in CS were therefore expected, yet serve as confirmation that our CS conditions induce a non-replicating state.

#### Arabinogalactan biosynthesis

The next layer of the cell wall is arabinogalactan (AG), which is covalently linked to both PG and mycolic acids (MA). AG is comprised of branched arabinose chains attached to galactan. The galactan chain is made by WbbL1 (MAB_3612c), GlfT1 (MAB_0202c), GlfT2 (MAB_0170), and likely other enzymes not yet annotated in *Mab*. Wbbl1 was only found in REP samples and Glft2 was significantly more abundant in REP than CS (4.8-fold, p = 1.4 × 10^−5^) (**Figure 4B**).

Since these enzymes are involved in early AG biosynthetic steps, it is possible that AG production is reduced in CS. In the periplasm, arabinose is added to the galactan portion by various arabinosyltransferases, including AftA-D. AftC and AftD were unchanged between CS and REP samples. The target of ethambutol, EmbC (MAB_0189c), was 1.8-fold more abundant in CS (p = 0.0029), while EmbA (MAB_0186c) and EmbB (MAB_0185c) were unchanged. DprE1 (MAB_0192c) and DprE2 (MAB_0191c) also play a role in AG biosynthesis; they were unchanged between CS and REP.

#### Mycomembrane biosynthesis

A distinguishing feature of the mycobacterial cell wall is the outer mycomembrane comprised of long-chain mycolic acids (MAs), including trehalose monomycolate and trehalose dimycolate. Many enzymes involved in MA biosynthesis, modification, and transport were less abundant in CS (**Figure 4C**). The key fatty acid synthesis enzyme Fas (MAB_1512) was 1.7-fold less abundant in CS (p = 0.00038). Next in the pathway, fatty acids are extended by the FAS-II complex of enzymes. Several FAS-II enzymes were less abundant in CS, including KasA (MAB_1877c; 2.5-fold), HadC (MAB_3896c; 2.3-fold), and MabA/FabG1 (MAB_2723c; 1.5-fold). Notably, a reduction in KasA protein levels is involved in biofilm formation in *Mycobacterium smegmatis*^55^. InhA (MAB_2722), a target of isoniazid, was unchanged between CS and REP. In *Mtb*, there are two pathways that make trehalose, OtsA/OtsB1 and TreX-TreY-TreZ^56^. We found that OtsA/OtsB1 were unchanged between CS and REP, while TreX (MAB_2690) was 1.8-fold (p = 0.013) more abundant in CS than REP. Taken together, these findings suggest that the as-yet uncharacterized TreX-TreY-TreZ pathway might play a more important role than OtsA/OtsB1 under nutrient-limited conditions in *Mab*. TreZ has not yet been identified in *Mab*.

Attachment of MA to AG is mediated by the antigen 85 complex (FbpA-D). We identified three of the four components of this complex as unchanged between CS and REP. The mycolic acid desaturase DesA1 has several putative orthologs in *Mab*, including MAB_3354, MAB_2550c, and MAB_0754c. Interestingly, two of these were significantly changed in opposite directions under CS. MAB_3354 was largely more abundant in CS (20-fold, p = 1.3 × 10^−9^, **Table S4**, excluded from heat map), while MAB_2550c was 1.9-fold more abundant in REP (p = 3.9 × 10^−5^). DesA2 (MAB_1237) was 1.4-fold less abundant in CS.

There were two mycomembrane proteins identified only in CS: MabR (MAB_1880c) and LpqY (MAB_1372). In *Mtb*, MabR (Rv2242) is a transcriptional repressor of the *fasII* operon^57^. Our proteomics data suggest that it could play a similar role in *Mab* because we observed several FAS-II enzymes were less abundant in CS. *Mab*’s LpqY (MAB_1372) has not been characterized, but in *Mtb* it is involved in sugar transport and trehalose recycling^58^. To summarize, our data suggest that *Mab* reduces MA biosynthesis in response to carbon starvation, which is similar to how *Mtb* responds to nutrient starvation^16,58,59^.

#### PIM biosynthesis

Phosphatidylinositol mannosides (**PIMs**) are abundant plasma membrane glycolipids in mycobacteria^60^. Most mycobacterial PIMs contain either two (PIM2) or six (PIM6) mannose residues (**Figure 5**)^61^. We examined the PIM biosynthetic machinery in our proteomics dataset and found that the key PIM acyltransferase, PatA (MAB_2895c), was 10.4-fold more abundant in REP than CS (p = 1.0 × 10^−7^) (**Figure 5B**). Other biosynthetic components were not found (PimE, PimB’) or unchanged (PimA). An uncharacterized, putative mannosyltransferase MAB_1122c was only found in REP samples. LpqW (MAB_1315), an essential lipoprotein, was more abundant in REP (3.5-fold; p = 0.00013). Studies of the ortholog from *M. smegmatis* suggest LpqW regulates the relative levels of PIMs versus lipoarabinomannans^62,63^.

**Figure 5.**
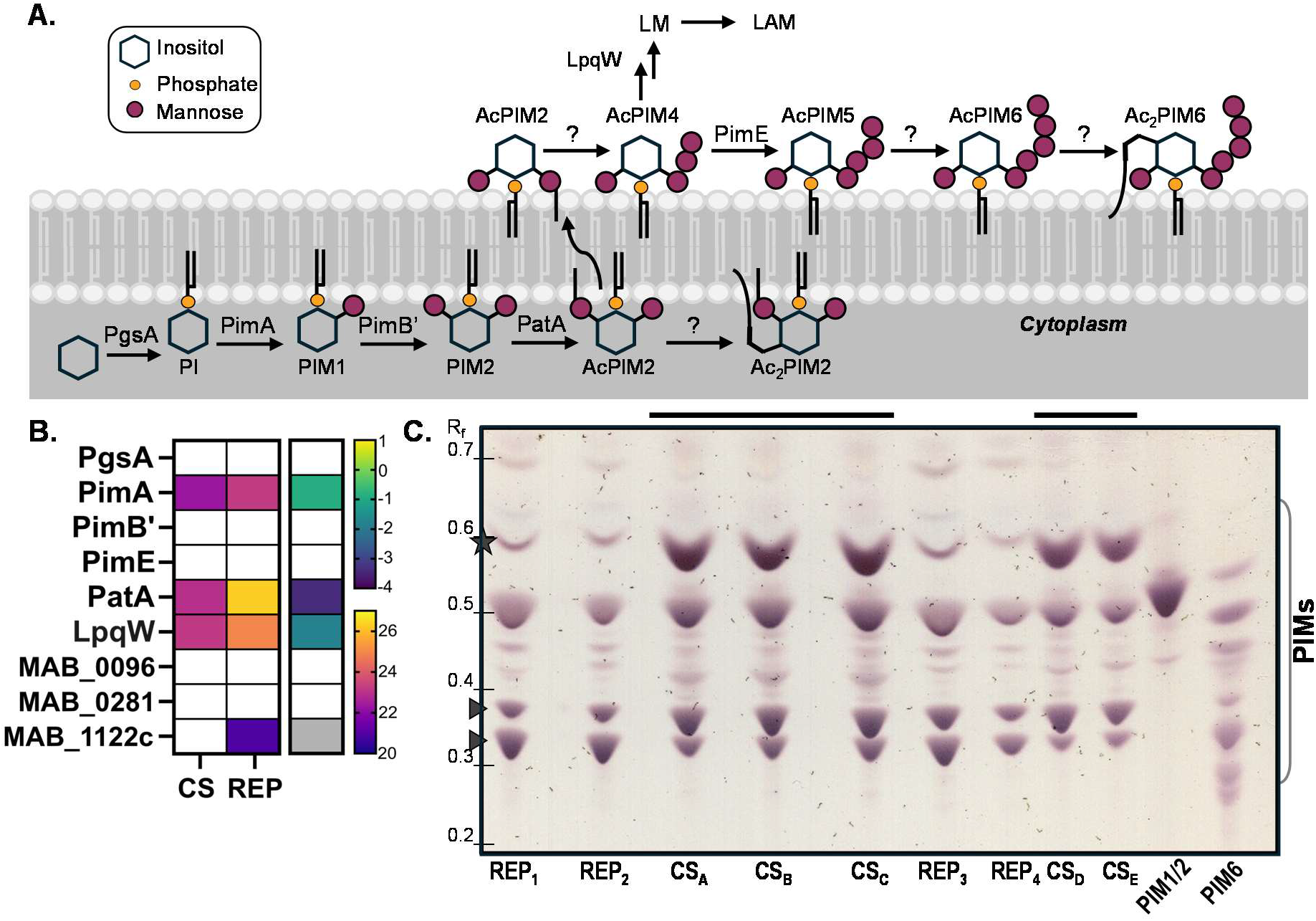
PIM biosynthesis is altered under CS. **A**. Schematic of the biosynthesis of mycobacterial PIMs. In *Mab*, the predominant PIMs are acylated PIM2 and PIM6. **B**. Heat map of mean log2 intensities (left map, warm scale) and corresponding log2 fold-change (right map, cool scale) in CS versus REP groups (n = 6) of proteins involved in PIM biosynthesis. A grey box indicates a protein was present in only one group (Group A: n ≥ 3, Group B: n ≤ 1). A white box indicates absence of value. Asterisks denote significance of the difference in presence (g-test) or mean intensities (ANOVA) between CS and REP (*: P ≤ 0.05, **: P ≤ 0.01, ***: P ≤ 0.001). **C**. TLC separation of normalized polar lipid extracts from cultures of *Mab*. PIMs were resolved in chloroform, methanol, ammonia, 0.1 M ammonium acetate, and water (180:140:9:9:23) and stained with orcinol. Putative PIMs are indicated by a bracket. The increased abundance of a PIM species at R_F_ ~ 0.6 is indicated with a star. Changes in relative abundance of two species (R_F_ = 0.3-0.4) are indicated by arrowheads. PIM1/2 and PIM6 purified from *M. tuberculosis* H37Rv were obtained from BEI Resources.

We sought to further describe the consequences of CS on PIMs. To do so, we analyzed the relative abundance of PIMs isolated from CS and REP cultures of *Mab*. Extracted polar lipids were resolved by thin layer chromatography (TLC) and stained with orcinol (**Figure 5C**). Loading was normalized for each sample based on the culture’s optical density. PIM1/2 and PIM6 isolated from *Mtb* were included in the analysis. We observed numerous resolved bands most likely corresponding to PIM2 and PIM6 species, based on their migration (i.e., retention factor or R_F_) and orcinol staining^61,64^. Notably, we observed an increased abundance in CS of a species just below R_F_ = 0.6. We have not identified the PIM species, but the migration suggests that it could be a di-acylated PIM2 (Ac_2_PIM2). This finding would be consistent with Morita and coworkers^61^ work, which demonstrated that PIM acylation is a mechanism used by mycobacteria, including *Mab*, to respond to stress. We additionally observed a change in relative intensity for two lower bands (R_F_ = 0.32 and 0.38), which likely correspond to a more mannosylated PIM species (i.e., PIM6).

#### Glycopeptidolipid biosynthesis

Glycopeptidolipids (GPLs) are abundant lipids on the outer cell envelope of some species of mycobacteria, including *Mab*^65^. *Mab* biosynthesizes both di-glycosylated GPL-2 and tri-glycosylated GPL-3 (**Figure 6**). In *Mab*, GPLs are linked to biofilm production, morphotype, and pathogenicity^66^. Generally, the presence of GPLs is associated with a “smooth” morphotype while an absence is associated with the “rough” morphotype. The rough morphotype is considered clinically relevant and is associated with late-stage, severe disease^67,68^. Genomic analysis of clinical isolates demonstrated that a rough morphology can result from mutations in genes found in the *gpl* locus, including *mps1, mps2, gap, mmpL4*, and *mmpS4*^68,69^. There is evidence that *Mab* can modulate GPL levels reversibly, although factors that cause these changes are largely unknown^70–72^. Prior work by Boutte and coworkers^73^ found that GPLs were unchanged by growth in rich Middlebrook 7H9 culture medium compared to either Hartmans-de Bont minimal medium or artificial cystic fibrosis sputum medium. In contrast, Jackson and coworkers observed an increase in the ratio of GPL-3 to GPL-2 and production of new GPL species in response to mucin exposure^72^, including in synthetic cystic fibrosis sputum medium and cystic fibrosis sputum.

**Figure 6.**
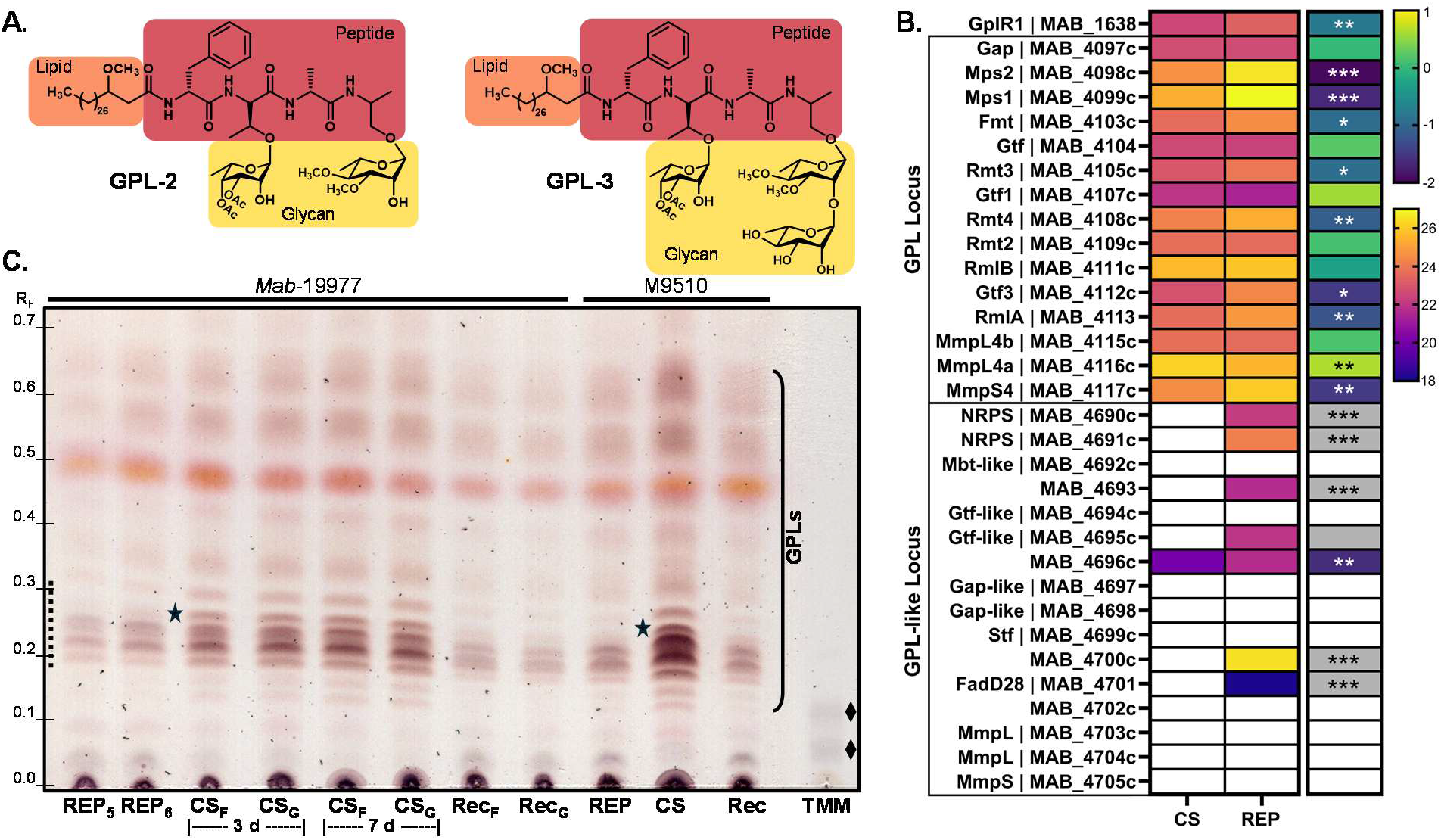
GPL biosynthesis is altered under CS. **A**. Structures of two GPLs made by *Mab*, GPL-2 and GPL-3. **B**. Heat map of mean log2 intensities (left map, warm scale) and corresponding log2 fold-change (right map, cool scale) in CS versus REP groups of proteins involved in GPL biosynthesis. A grey box indicates a protein was present in only one group (Group A: n ≥ 3, Group B: n ≤ 1). A white box indicates absence of value. Asterisks denote significance of the difference in presence (g-test) or mean intensities (ANOVA) between CS and REP (*: P ≤ 0.05, **: P ≤ 0.01, ***: P ≤ 0.001). NRPS = non-ribosomal peptide synthase. **C**. TLC analysis of normalized total lipid extracts from *Mab* 19977 and a clinical isolate, M9510. Total lipids were resolved in chloroform/methanol/water [90:10:1, v:v:v] and glycolipids (e.g., GPLs) were visualized by orcinol staining. Putative GPLs are indicated by a bracket, with the region (R_F_ = ~0.2-0.3) marked by a dashed line predicted to correspond to GPL-3. Black stars (R_F_ = 0.26) indicate a species detected only in CS. Trehalose monomycolate (TMM) purified from *M. tuberculosis* H37Rv was obtained from BEI Resources and included as a reference marker (diamonds).

Factors that dynamically impact GPL production in response to environment are unknown. We therefore analyzed our proteomics data for changes associated with CS (**Figure 6B)**. First, we found that the transcription factor GplR1 (MAB_1638) was 1.7-fold more abundant in REP than CS (p = 0.0097). Recent work demonstrated that the GplR1 is required for GPL production^74^ and influences the expression of both the *gpl* locus (MAB_4097c—MAB_4117c) and the *gpl*-like locus (MAB_4689c—MAB_4705c). We found that most of the proteins encoded in these two loci were more abundant in REP than CS. This evidence supports that CS impacts GPL biosynthesis.

Based on our proteomics data, we hypothesized that the GPL content would change between CS and REP cultures. We compared total lipid extracts (**Figure 6C**) and polar lipid extracts (**Figure S2**) isolated from REP and CS cultures of *Mab*. We additionally collected lipids from recovered (Rec) *Mab*, which was obtained by diluting and culturing carbon-starved *Mab* for one day in complete, nutrient-rich medium. Normalized extracts were analyzed by TLC, and we observed higher overall levels of GPLs in CS. A set of bands that migrate and stain like GPL3^75,76^ were particularly abundant at both 3 and 7 days of carbon starvation (**Figure 6C**). We also observed a species associated with CS (R_F_ = 0.26). The changes in GPL content were reversible, with reversion to a profile similar to REP in Rec samples. We additionally analyzed total lipids isolated from a clinical strain, *Mab* M9510 (subspecies *massiliense*)^77^. We observed similar changes in GPLs in this strain cultured under CS, suggesting that this response to CS is conserved across subspecies. Overall, our results suggest that CS results in a reversible enrichment of GPLs in *Mab*, a novel finding that warrants further exploration.

### Energy Metabolism

#### Oxidative Phosphorylation

Mycobacteria are obligate aerobes with the ability to energetically adapt to variable environments and to sustain metabolism in the absence of growth^78^. Unsurprisingly, most proteins involved in oxidative phosphorylation were unchanged in CS (**Figure 7A**). Of the differentially abundant proteins, most were moderately more abundant in CS compared to REP. The most prominent complex-wide difference was in the type I NADH dehydrogenase (NDH-1, complex 1). This complex is the more energy efficient of two NADH dehydrogenases in *Mab*^78^. Five of the identified NDH-1 proteins were significantly more abundant in CS, including NuoG (1.7-fold, p = 0.027), NuoH (3.5-fold, p = 0.017), NuoM (2.2-fold, p = 0.0046), and two were only identified in CS (NuoF and NuoN, p <0.006). No NDH-1 proteins were more abundant in REP compared to CS. The overall increase in NDH-1 subunits under CS suggests that *Mab* implements a mechanism of energy conservation under CS. This result agrees with previous reports of transcriptional upregulation of the NDH-1 operon in *Mab* in response to artificial sputum^12^ and in *M. smegmatis* under carbon-limited growth^79^. We also previously identified similar increased abundance of NDH-1 proteins in *Mtb* grown under CS^16^.

**Figure 7.**
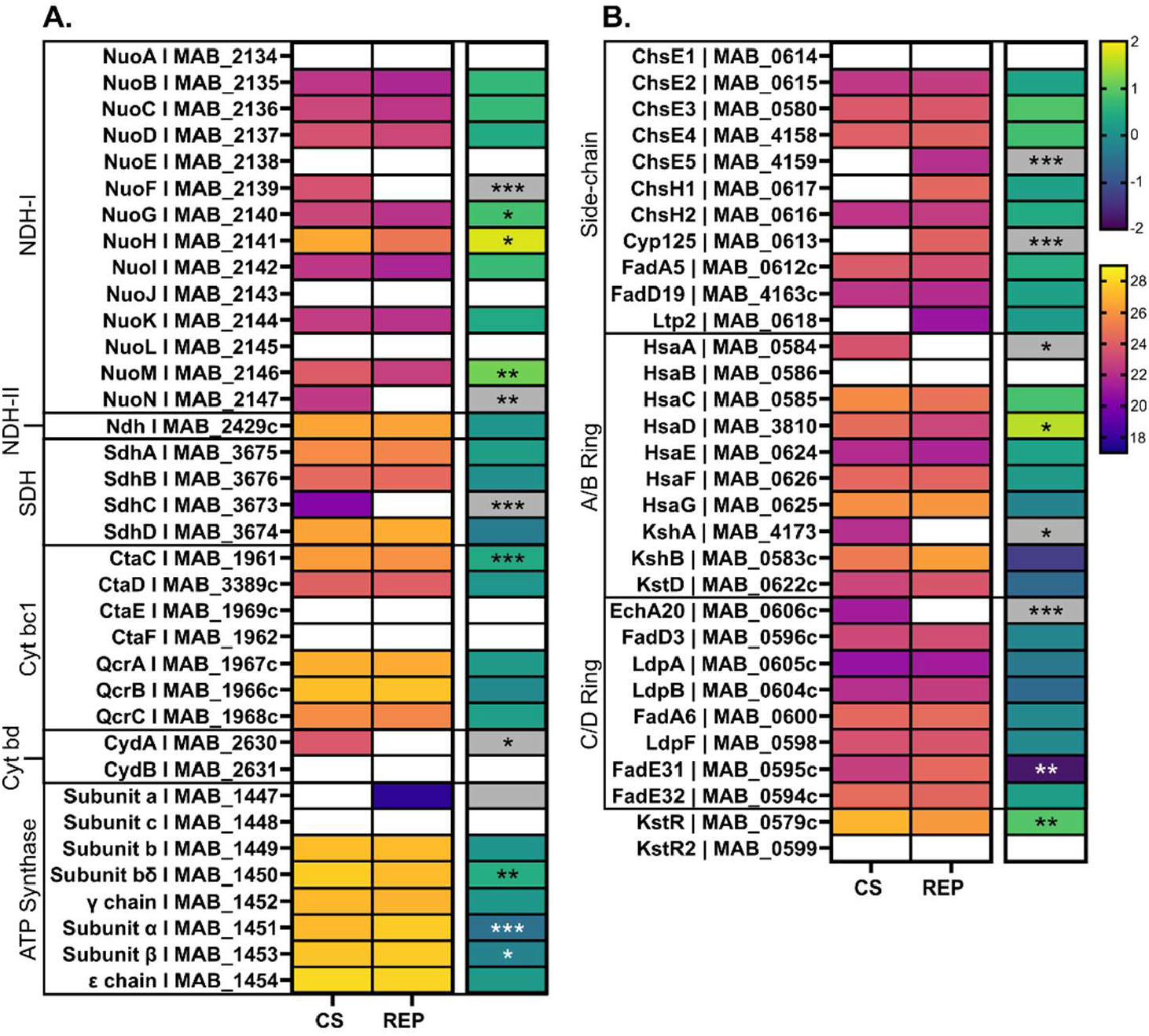
Regulation of energy and cholesterol metabolism under CS. Heat maps of mean log2 intensities (left map, warm scale) and corresponding log2 fold-change (right map, cool scale) in CS versus REP groups (n = 6) of proteins involved in oxidative phosphorylation (**A**) and cholesterol catabolism (**B**). Scale bars are applicable across all corresponding maps. A grey box indicates a protein was present in only one group (Group A: n ≥ 3, Group B: n ≤ 1). A white box indicates absence of value. Asterisks denote significance of the difference in presence (g-test) or mean intensities (ANOVA) between CS and REP (*: P ≤ 0.05, **: P ≤ 0.01, ***: P ≤ 0.001).

Several established anti-mycobacterial drugs target oxidative phosphorylation and energy production remains an attractive target for novel drug development. The targets of clofazimine, type 2 NADH dehydrogenase (Ndh, MAB_2429c)^80^, and bedaquiline, ATP synthase subunit c (AtpE, MAB_1448)^81^, were unchanged in CS relative to REP (**Table S4**). The anti-TB compound GaMF1 targets the γ subunit (AtpG, MAB_1452)^82^, which was also unchanged with CS.

#### Cholesterol Catabolism

Cholesterol is a primary source of energy for mycobacteria during infection^83^. Specifically in *Mab*, genes involved in cholesterol catabolism were upregulated in a macrophage infection model^31^ and a cholesterol catabolism mutant strain (Δ*hsaD*) was unable to replicate in macrophages^84^. These findings suggest that *Mab* depends on the ability to break down cholesterol to survive intracellularly. We observed notable changes in cholesterol catabolism in response to CS (**Figure 7B**). While two enzymes involved in side-chain degradation were less abundant in CS, all A/B ring cleavage enzymes that were significantly different were more abundant in CS. This included HsaD, whose gene is located separate from the *hsaACB* operon uniquely in *Mab*^84^. Interestingly, KstR, the TetR family repressor that regulates transcription of both side-chain and A/B Ring degradation enzymes^85^, was also more abundant in CS (1.9-fold, p = 0.01).

### Sulfate metabolism

Mycobacteria acquire sulfate from the environment to produce a variety of sulfated metabolites^86,87^ (**Figure 8**). Many of the key enzymes in sulfate metabolism have been characterized in *Mtb* and are conserved in *Mab*. We examined these sulfate metabolic enzymes within our proteomic data to identify changes between CS and REP (**Figure 8B**). A multi-protein sulfate importer, named SubI-CysT/W/A1, is used to acquire sulfate from the environment. We identified two of the components, CysA1 (MAB_1655), which was 1.2-fold more abundant in REP than CS, and CysT (MAB_1653), which was only identified in CS. Several enzymes in the sulfate assimilation pathway were more abundant in REP, including CysC (MAB_1544c), CysH (MAB_1661c), and the phosphatase CysQ (MAB_2115). The sulfotransferase MAB_1546c is an ortholog of the *Mtb* enzyme Stf0 (Rv0295c, 74.9% sequence identity), which sulfates trehalose^88^. MAB_1546c was 2-fold more abundant in REP than CS (p = 0.040). The hydrolysis of sulfates is catalyzed by sulfatases, which also changed in response to CS. Specifically, AtsG (MAB_1547c) was identified only in REP and AtsF (MAB_4469c) was identified only in CS. To summarize, these findings suggest that various changes in sulfate metabolism occur in CS.

**Figure 8.**
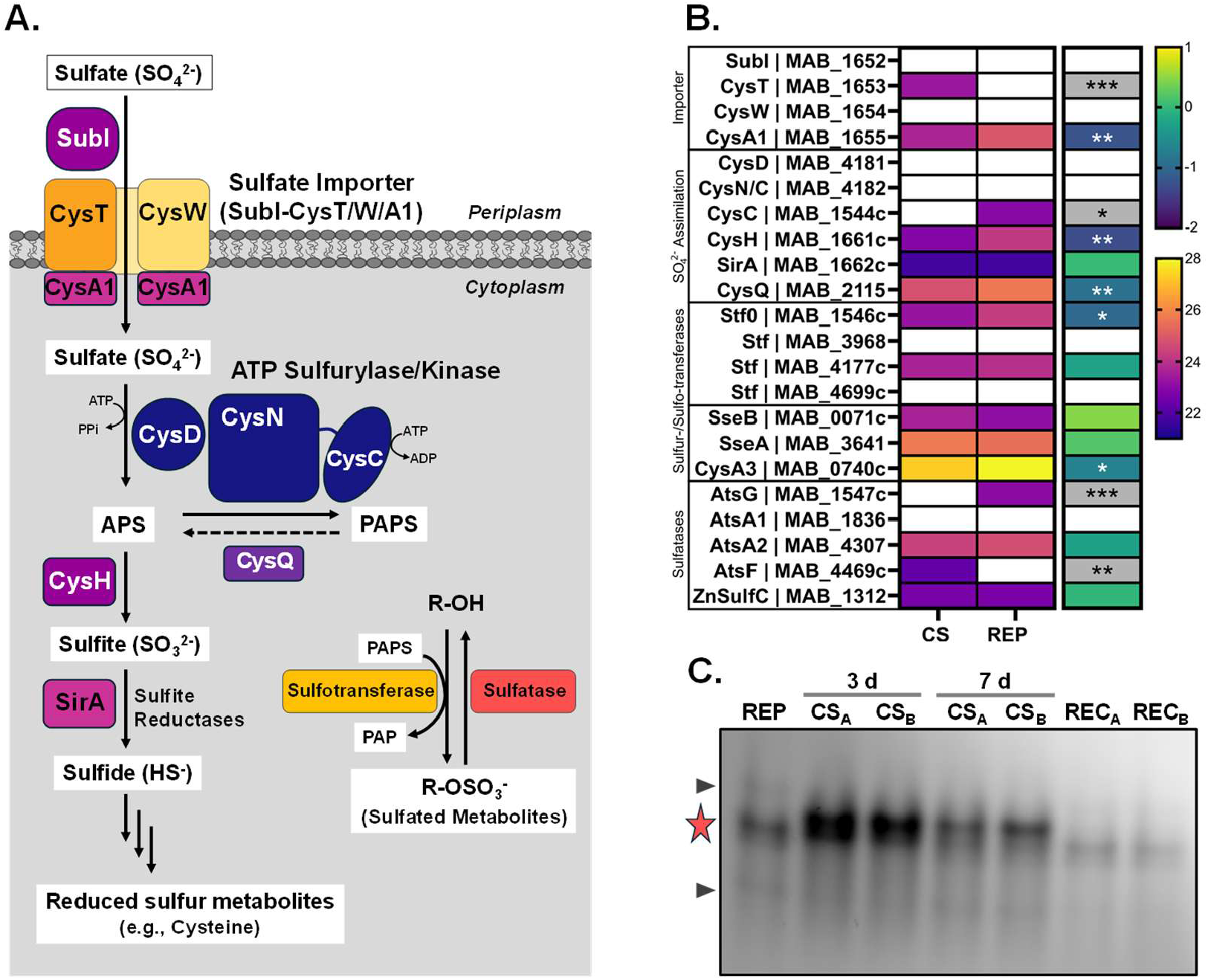
Sulfate metabolism is changed in carbon starved *Mab*. **A**. The sulfate assimilation pathway. Inorganic sulfate is acquired through the sulfate importer. It is activated to adenosine-5’-phosphosulfate (APS) by CysD/CysN/C and converted to 3’-phosphoadenosine-5’-phosphosulfate (PAPS), the universal sulfate donor. PAPS is used by sulfotransferases to sulfate biomolecules. The reverse reaction is catalyzed by sulfatases. Separately, in the reductive path, APS can be converted to sulfite and sulfide. **B**. Heat map of mean log2 intensities (left map, warm scale) and corresponding log2 fold-change (right map, cool scale) of sulfate metabolism-involved proteins in CS versus REP groups (n = 6). A grey box indicates a protein was present in only one group (Group A: n ≥ 3, Group B: n ≤ 1). A white box indicates absence of value. Asterisks denote significance of the difference in presence (g-test) or mean intensities (ANOVA) between CS and REP (*: P ≤ 0.05, **: P ≤ 0.01, ***: P ≤ 0.001). **C**. Sulfatase activity detected in gel-resolved lysates of *Mab* incubated with DDAO-sulfate. The star highlights sulfatase activity induced in CS. Other sulfatase activity indicated with arrowheads. REC = recovered from CS.

We did a preliminary assessment of *Mab* sulfatase activity in REP and CS conditions. We used a method for detecting active sulfatases in native protein gels using a fluorogenic sulfatase probe, DDAO-sulfate, as described previously^89^. Briefly, normalized total lysates were resolved by native gel electrophoresis, incubated with DDAO-sulfate, and imaged. We observed an increase in sulfatase activity in CS samples (**Figure 8C**). Upon recovery from CS (REC), activity reverted to lower levels observed in REP. We have not identified the enzymes associated with each band, but we speculate that the CS-induced fluorescent band is most likely AtsF because this sulfatase was only identified in CS samples.

### Cobalamin biosynthesis

The cofactor Cobalamin, also known as Vitamin B12, has a tetrapyrrole framework that chelates a central cobalt ion (**Figure 9**). This is a biosynthetically expensive molecule, requiring 30 reactions to make. Many mycobacterial pathogens have the necessary set of enzymes for *de novo* cobalamin biosynthesis^90,91^ and cobalamin has been found in several strains of *Mab*^90^. Our data indicate that cobalamin biosynthesis is reduced in CS (**Figure 9B**). Specifically, several of the enzymes in the pathway were found only in REP: CobD (MAB_1898), CobF (MAB_1037), CobK (MAB_2197), and CobL (MAB_2195). Two components of the Uroporphyrinogen III pathway were >2.5-fold more abundant in REP (CysH and HemD), while HemY (MAB_2985c) was 2.0-fold more abundant in CS (p = 0.026). The cobalt transporter sub-unit CbtA (MAB_0551) was 8.7-fold more abundant in REP (p = 0.0038). Two cobalamin-dependent enzymes, MetH (MAB_2129) and MutB (MAB_2711c) were unchanged between REP and CS. Overall, our data indicate that the biosynthetic machinery to make cobalamin is present in both conditions, but more abundant in REP than CS.

**Figure 9.**
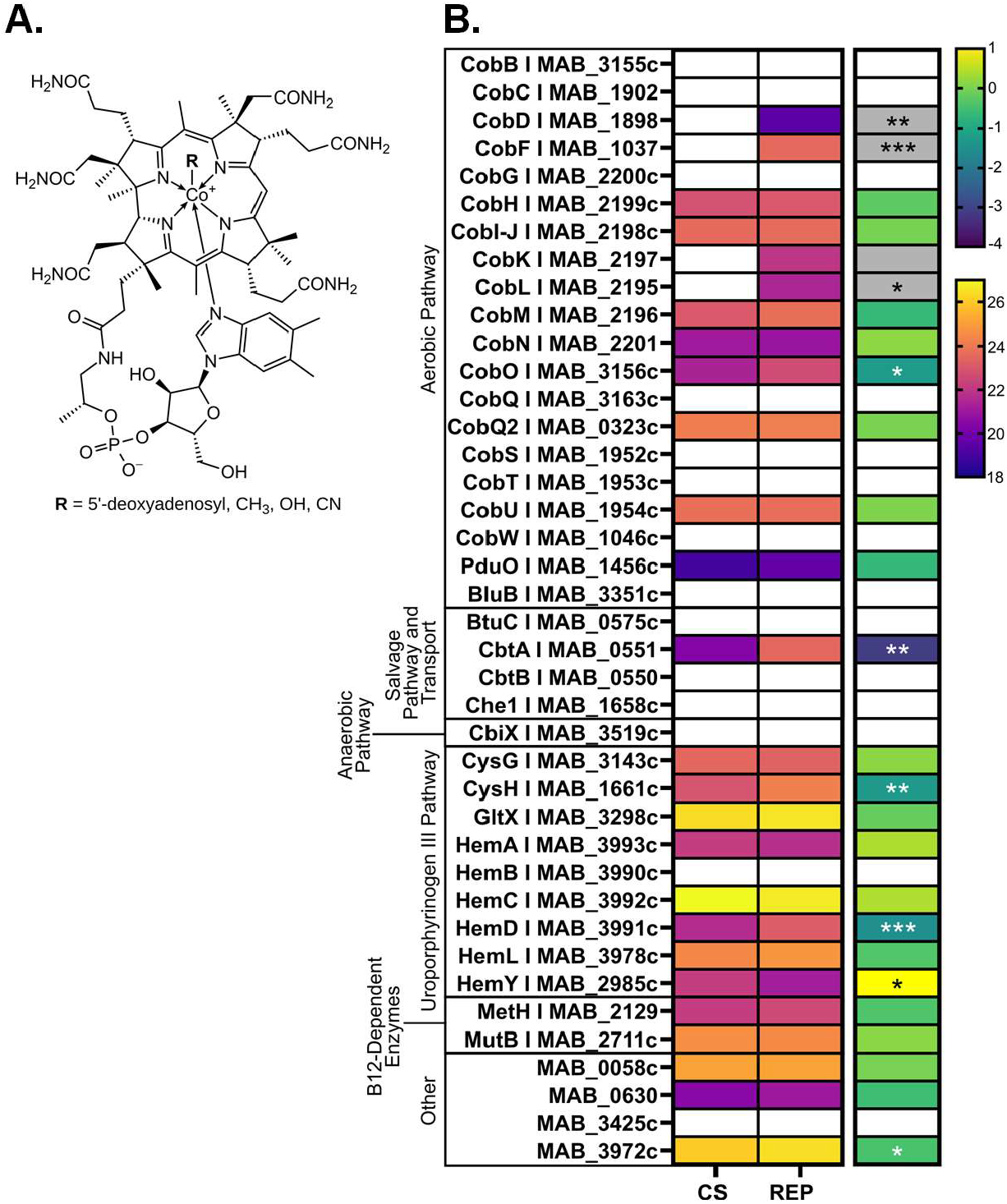
Regulation of *Mab* cobalamin biosynthesis under CS. **A**. Structure of cobalamin (Wikipedia). **B**. Heat map of mean log2 intensities (left map, warm scale) and corresponding log2 fold-change (right map, cool scale) of cobalamin biosynthesis in CS versus REP groups (n = 6). A grey box indicates a protein was present in only one group (Group A: n ≥ 3, Group B: n ≤ 1). A white box indicates absence of value. Asterisks denote significance of the difference in presence (g-test) or mean intensities (ANOVA) between CS and REP (*: P ≤ 0.05, **: P ≤ 0.01, ***: P ≤ 0.001).

### Iron acquisition, iron-sulfur proteins, and cytochrome P450s

Mycobacteria require iron for growth and infection. The base broth used in both our CS and REP media contains trace metals, including ferric ammonium citrate. Iron acquisition did not appear to be impacted by CS (see **Table S4**). In contrast, notable differences in iron-sulfur [Fe-S] cofactor biosynthesis, [Fe-S] proteins, and cytochrome P450s were found in CS compared to REP cultures.

### [Fe-S] cluster biosynthesis

Mycobacteria use the sulfur utilization factor (SUF) machinery to make [Fe-S] clusters^92^. The *Mab* SUF operon includes seven genes: sufRBDCSUT (MAB_2750c − MAB_2744c). SufU (MAB_2745c) was found only in REP, while four other components were more abundant in REP than CS: SufB, SufD, SufC, SufS (**Figure 10**). The [Fe-S] cluster delivery protein SufA (MAB_1957) was more abundant in REP (1.6-fold, p = 0.013) while two others, Mrp (MAB_1366c) and SufT (MAB_2744c), were unchanged. Our findings suggest that [Fe-S] cluster biosynthesis is down-regulated in response to carbon starvation.

**Figure 10.**
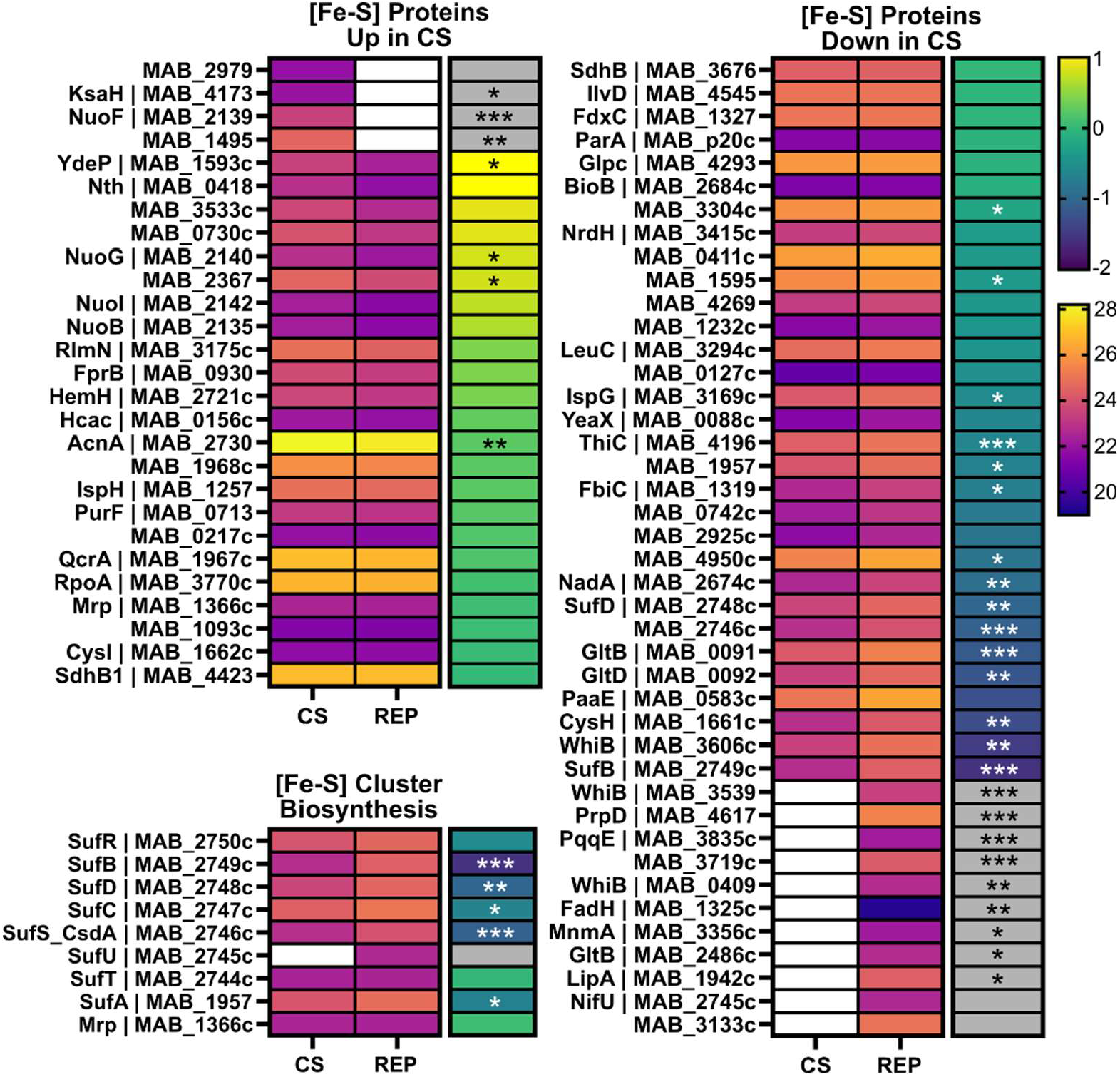
Regulation of *Mab* [Fe-S] cluster biosynthesis enzymes and [Fe-S] proteins under CS. Heat maps of mean log2 intensities (left maps, warm scale) and corresponding log2 fold-change (right maps, cool scale) in CS versus REP groups (n = 6). Scale bars are applicable across all corresponding maps. A grey box indicates a protein was present in only one group (Group A: n ≥ 3, Group B: n ≤ 1). A white box indicates absence of value. Asterisks denote significance of the difference in presence (g-test) or mean intensities (ANOVA) between CS and REP (*: P ≤ 0.05, **: P ≤ 0.01, ***: P ≤ 0.001).

### [Fe-S] cluster proteins

A recent review summarized the roles of mycobacterial [Fe-S] cluster proteins in diverse processes including virulence, persistence, metabolism, and antibiotic resistance^92^. The authors identified 126 [Fe-S] cluster proteins encoded by *Mab*; 69 of these were identified in our proteomic samples. Our data indicate that *Mab*’s [Fe-S] cluster proteins are differentially expressed in response to CS. Specifically, eleven [Fe-S] cluster proteins were found only in REP samples, including two WhiB transcriptional regulators (MAB_3539 and MAB_0409). A third WhiB regulator, MAB_3606, was 2.8-fold more abundant in REP (p = 0.0017). On the other hand, four [Fe-S] cluster proteins were only found in CS (i.e., KsaH, NuoF, MAB_1495, and MsrB). KsaH (MAB_4173) is involved in cholesterol metabolism and NuoF (MAB_2139) is involved in energy production (see **Figure 7**). MAB_1495 is a probable oxidoreductase. Lastly, MsrB (MAB_2979) is a peptide-methionine (R)-S-oxide reductase.

### Cytochrome P450s

Cytochrome P450s (CYPs) are heme-dependent monooxygenases and potential drug targets. The genome of *Mab* encodes 25^93^, which is >250-fold greater “CYP density” than the human genome^94^. A complete list of *Mab* CYPs are provided in the Supporting Information (**Table S4**) and were named according to subfamily, as described^95^. We identified eight CYPs in our proteomic samples (**Figure 11**). The best described is CYP51, an ancient CYP found across species and kingdoms. In *Mtb*, CYP51 (Rv0764c) is a sterol demethylase^96^. The corresponding CYP51 in *Mab* (MAB_1214c) was identified only in REP samples. In our dataset, four additional CYPs were identified only in REP samples: CYP164A (MAB_0276), Cyp125A (MAB_0613), CYP1110B1 (MAB_0917c), and CYP135B1 (MAB_4693). *Mtb*’s CYP125A (Rv3545c) plays a key role in cholesterol metabolism and survival *in vivo*^97,98^, while CYP135B1 may detoxify the anti-TB drug SQ109^99^. The functional consequence of the downregulation of these enzymes under CS in *Mab* is unclear. We identified three CYPs that were not significantly changed between REP and CS, namely Cyp132 (MAB_1426), Cyp108b (MAB_3825), and Cyp140B (MAB_4457). We identified nine putative redox partners (e.g., ferredoxins), and all but one were unchanged between REP and CS (**Figure 11B**).

**Figure 11.**
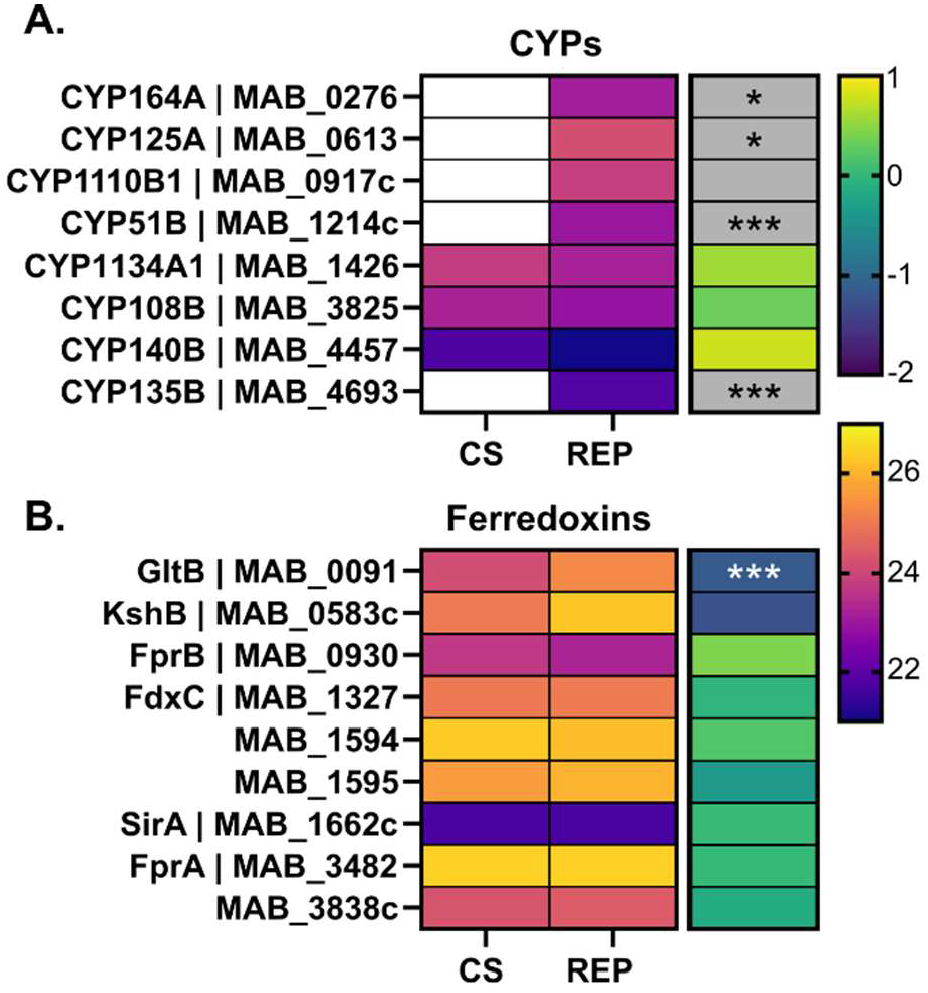
Regulation of *Mab* cytochrome P450s and ferredoxins under CS. Heat map of mean log2 intensities (left map, warm scale) and corresponding log2 fold-change (right map, cool scale) of CYPs (**A**.) and ferredoxins (**B**.) in CS versus REP groups (n = 6). Scale bars are applicable across all corresponding maps. A grey box indicates a protein was present in only one group (Group A: n ≥ 3, Group B: n ≤ 1). A white box indicates absence of value. Asterisks denote significance of the difference in presence (g-test) or mean intensities (ANOVA) between CS and REP (*: P ≤ 0.05, **: P ≤ 0.01, ***: P ≤ 0.001).

## Conclusions

The ability to enter and maintain a non-replicating state is critical for *Mab* to maintain chronic infections in humans. It is therefore crucial to understand this physiological state. Several *in vitro* models mimicking microenvironmental conditions within the host have been developed that induce a non-replicating state in *Mab*. We recently described a simple and accessible model based on carbon starvation, which induces a non-replicating but viable state^19^. Herein, we provide the first detailed analysis of the proteomic changes associated with CS, broadening our understanding of persistent *Mab*.

We found that *Mab* uses two-component signal transduction systems to respond to CS. Similar to the hypoxic response^11,30^, the stress response regulators DosRS and many corresponding downstream proteins were more abundant under CS. We also found widespread alterations in enzymes responsible for secretion, nutrient biosynthesis, acquisition, and metabolism. Consistent with other mycobacterial persistence models, we observed a shift toward energy conservation within oxidative phosphorylation under CS, with an increased abundance of the energy efficient type I NADH dehydrogenase complex.

Our data support that the non-replicating state is maintained by altering the cell envelope, including PG biosynthesis and degradation. Numerous changes in the cell envelope machinery were noted, including significant changes in the GPL locus. We confirmed the effect of these proteomic changes by demonstrating that two classes of glycolipids, PIMs and GPLs, underwent substantial changes in CS. The outcome of these changes is unclear but could be profound considering the cell envelope is both a barrier and the first point of contact with the host.

Overall, the detailed description of *Mab*’s proteomic remodeling in CS broadens our understanding of the molecular mechanisms and characteristics of *Mab* non-replicating persistence. The data provided here will serve as a rich resource for further work using *in vitro* persistence models.

## Supplemental Material

Detailed “Materials and Methods”

Figures S1 and S2

Excel Files: Table S1 – Table S4

## Data Availability

Proteomics data have been deposited in MassIVE (see Cover Letter for details).

## Acknowledgements

Funding for the Beatty group was provided by the National Institute of Health (NIAID: R01 AI149737 and NIGMS: R35 GM149559). GL was funded by NIH R01 AI155664. PNNL is operated by Battelle for the DOE under contract DE-AC05-76RL01830. We are grateful to Berit Blume, Victoria Halls, and Carsten Schultz (OHSU) for their expertise in lipid extraction and TLC analysis. We thank Priscila Lalli for mass spectrometry and Matt Monroe (PNNL) for proteomics data support. The following reagents were obtained through BEI Resources, NIAID, NIH: *Mycobacterium tuberculosis*, Strain H37Rv, purified trehalose monomycolate (TMM; NR-48784), trehalose dimycolate (TDM, NR-14844), phosphatidylinositol mannoside 1/2 (PIM1,2; NR-14846), and PIM6 (NR-14847).

